# Metabolites cross the membrane, the transcriptome does not follow: co-localization is required for the cyanobacterial response to a yeast partner

**DOI:** 10.64898/2026.11.05.764704

**Authors:** Pavlo Bohutskyi, Zachary Johnson, Marija Veličković, Young-Mo Kim, Margaret S. Cheung

## Abstract

Interactions between phototrophs and heterotrophs are usually explained by cross-feeding, yet whether that exchange alone accounts for the phototroph’s response has rarely been tested. We grew the sucrose-secreting cyanobacterium *Synechococcus elongatus* with the yeast *Rhodotorula toruloides* as a synthetic community, either in mixed co-culture or in membrane-separated co-culture (0.22 µm pore size), each with a matched axenic control. Metabolite exchange occurred in both configurations: sucrose reached the yeast and was consumed, and yeast-derived metabolites reached the cyanobacterium. The yeast reached a higher final density behind the membrane, whereas the cyanobacterium gained about 9% over its axenic control only in mixed co-culture. Yet the cyanobacterial transcriptome responded only with co-localization: expression of 51 genes changed consistently across time points (day-averaged q < 0.05), against one gene behind the membrane, and the number of differentially expressed genes rose from 20 on day 4 to 91 and 127 on days 6 and 8. Nine regulatory programs (iModulons), mainly nitrogen and photosynthesis, responded in mixed co-culture and none behind the membrane. Reactor configuration alone changed 40 genes in axenic culture. These results suggest that co-localization, in addition to metabolite exchange, was required for the heterotrophic partner to trigger a response in the phototroph. Co-culture studies that infer interactions from shared metabolites may miss effects that depend on spatial structure.

**Importance:** Partnerships between light-powered microbes and the heterotrophs living alongside them move large amounts of carbon in oceans and soils, and are now built on purpose to turn carbon dioxide into fuels and chemicals. Such partnerships are usually described by the chemicals the partners trade, so we asked whether those chemicals are enough. The cyanobacterium *Synechococcus elongatus* and the yeast *Rhodotorula toruloides* were grown together in one vessel, or in two chambers divided by a membrane that molecules cross but cells cannot. The yeast fed on cyanobacterial sugar in both, yet the cyanobacterium changed its gene activity only when the two shared a space. Partners in nature are usually clustered rather than evenly mixed, so closeness may contribute as much as the chemicals themselves. Keeping partners apart in engineered cultures can still deliver carbon to the heterotroph, but may not deliver the benefits of partnership to the phototroph.

## Introduction

Associations between phototrophs and heterotrophs are widespread in aquatic and terrestrial ecosystems, from the phycosphere of planktonic algae to microbial mats and lichens, and they contribute to primary production and nutrient cycling (1–3). In these associations, the phototroph releases fixed organic carbon, and the heterotroph provides functions that the phototroph cannot perform alone (3, 4). Heterotrophic partners recycle nutrients and support long-term stability of the community (5, 6), supply essential vitamins (4, 7, 8), and relieve oxidative stress, which improves phototroph viability (9–11). These functions are applied in synthetic phototroph-heterotroph consortia for light-driven bioproduction, in which heterotrophic partners improve growth, stability and product yield (12, 13). Major cross-species interactions and the benefits to the phototroph are generally attributed to metabolites exchanged between the partners (1, 14, 15), yet whether exchange alone accounts for the phototroph response has received little attention.

Synthetic binary consortia allow these exchanges, and their effect on the phototroph, to be studied under defined conditions. Heterotrophic bacteria and yeasts benefit sucrose-secreting *Synechococcus elongatus* PCC 7942 (11, 16, 17), the same *cscB*/SPS strain exchanges fixed carbon for fixed nitrogen with *Azotobacter vinelandii* (18), and *Alteromonas* alters the growth, decline and transcriptome of *Prochlorococcus* (19, 20). Community metabolic models link such partners through a shared metabolite pool and explain the benefits by exchange fluxes within it (14, 21). In natural communities, however, the distance between partners also matters. Many phytoplankton-bacteria exchanges take place in the phycosphere, the microenvironment immediately surrounding the algal cell, and depend on close proximity (2); bacteria attach to algal cells and form a joint extracellular matrix with them (22). In hot spring mats, *Synechococcus* and *Roseiflexus* are proposed to exchange glycolate and lactate (23), and in lichens, fungal and algal cells form specialized contacts thought to channel solutes between the partners (24). In engineered consortia, algae associate with filamentous fungi (25), and spatial structure determines whether a community persists (26). Membrane-separated co-cultures, which allow metabolite exchange but prevent cell contact, provide a direct test of what exchange alone contributes (27, 28). Across a membrane, the phototroph still supports the growth of its heterotrophic partner, and membrane-separated cultures of *S. elongatus* and *R. toruloides* revealed an exchange of sugars for sugar alcohols that mirrors natural cyanolichens (28, 29). Whether, and to what extent, this exchange alone accounts for the benefit returned to the phototroph, and whether its molecular response and phenotypic adjustment are the same with and without co-localization, have not been examined.

Molecular responses of the phototroph to its partner have so far been characterized mainly in shared-volume co-cultures. Transcriptomes, proteomes and metabolomes of *S. elongatus* differ between co-culture and monoculture (12, 14, 17, 18), although these were measured mostly at a single growth stage and the interactions change over time (19, 20). The magnitude varies between systems: in dual transcriptomes of microalgae with phycosphere bacteria, the algal transcriptomes were reported to change little (30). Comparing spatial configurations adds a further difficulty, because separating the partners also changes the reactor, and physicochemical factors such as mixing, gas exchange, pH and buffering capacity alter the outcome of algae-bacteria interactions (31). A test of the shared-pool view therefore requires a matched control for each configuration and repeated sampling. We hypothesized that, if exchange through a shared metabolite pool were sufficient, *S. elongatus* would respond in the same way whether its partner shared the culture volume or was separated by a membrane.

Here we asked whether metabolite exchange alone accounts for how a phototroph responds to a heterotrophic partner, using *S. elongatus* and the yeast *Rhodotorula toruloides*, a pairing in which the cyanobacterium supplies sucrose to the yeast. A 0.22 µm membrane separates the partners while leaving the diffusible pool shared, so comparing a membrane-separated co-culture with a mixed one distinguishes what reaches the phototroph by diffusion from what requires proximity. Each configuration carried its own axenic *S. elongatus* control, separating the effect of the partner from that of the reactor configuration. Following both configurations over ten days, we tracked growth and the exometabolome and resolved the response of *S. elongatus* at the level of genes and of regulatory programs. For the latter we used iModulons, sets of co-regulated genes learned from a 407-sample transcriptome compendium, whose activities reflect the activity of their regulators (32, 33). Metabolites crossed the membrane in both directions, yet *S. elongatus* reorganized its gene expression only when the partners shared a volume and could interact directly: 51 genes and nine iModulons responded in the mixed reactor, compared with one gene and no iModulons when *R. toruloides* was separated by the membrane. These results suggest that co-localization, and not metabolite exchange alone, shapes the response of the phototroph to its heterotrophic partner.

## Results

### Growth, sucrose exchange, and the separation of partner and reactor effects

To separate the effect of the partner from that of the reactor configuration, each configuration was paired with an axenic *S. elongatus* control in the same reactor type (**Fig. 1a**). This design defines four comparisons, each expressed as the first group relative to the second: co-culture against axenic culture in the mixed reactor (mixed CC vs Ax; CC, co- culture; Ax, axenic) and in the *S. elongatus* (Se) chamber of the two-chamber membrane bioreactor (mBR Se CC vs Ax), the *R. toruloides* (Rt) chamber against the cell-free chamber (mBR Rt chamber), and the two axenic cultures against each other (mBR Ax vs mixed Ax), a configuration control. Rt was added at 44.65 h, after the day 2 sampling, so days 1 and 2 provide a baseline in each comparison.

**Figure 1.**
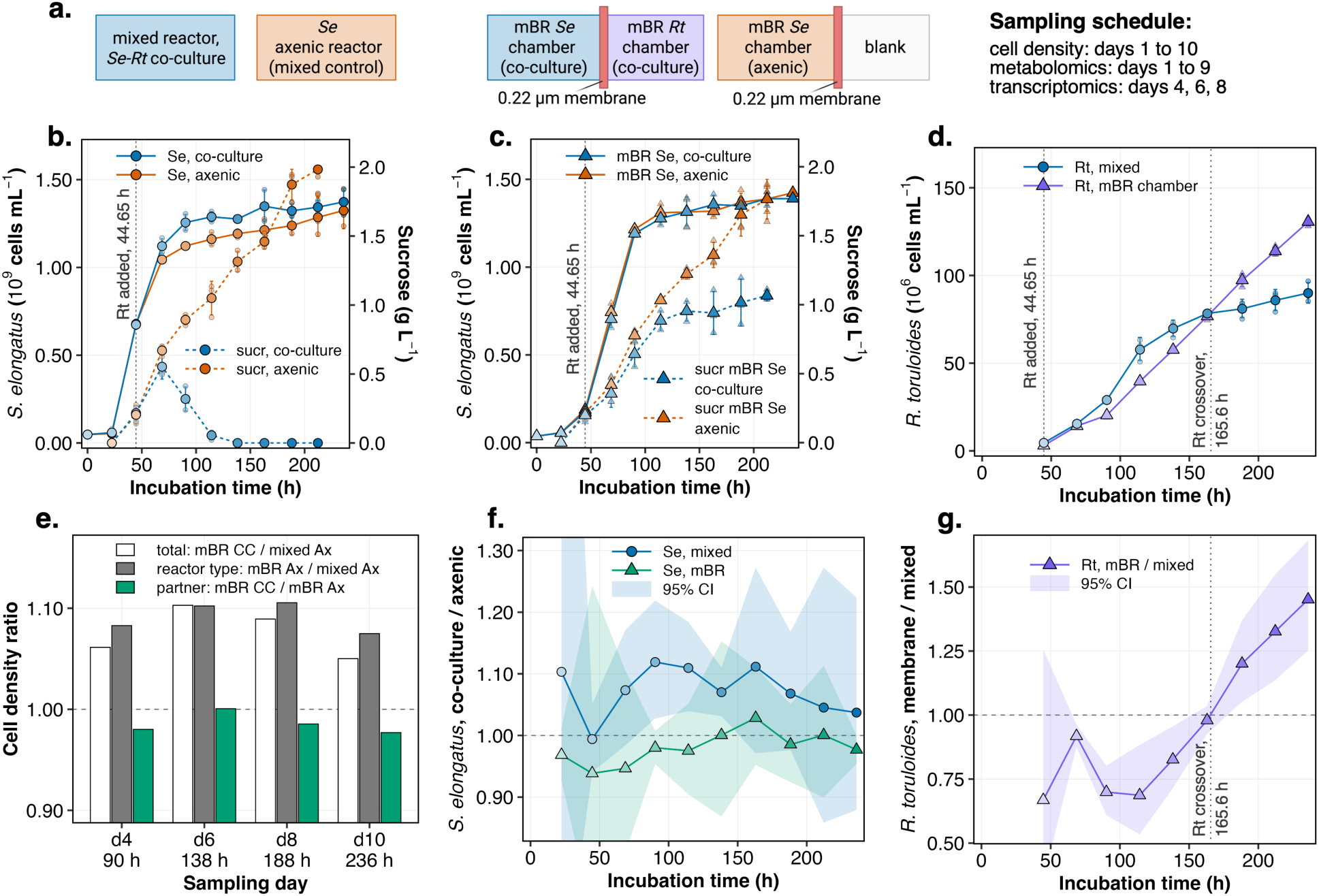
R. toruloides raised S. elongatus density only in the mixed reactor, whereas the higher density in the mBR reflected the configuration. (a) A mixed reactor (one shared volume) and a two-chamber mBR (0.22 µm membrane between the partners), each with its own axenic Se control; in the axenic mBR the second chamber holds medium alone. (b, c) Se cell density (solid, left axis) and extracellular sucrose (dashed, right axis) in CC and Ax, for the mixed reactor (b) and the mBR Se chamber (c). (d) Rt cell density in both configurations. (e) The previously used comparison, mBR CC vs mixed Ax, separated into a reactor-type part (mBR Ax over mixed Ax) and a partner part (mBR CC over mBR Ax). (f) Se density in CC relative to its own Ax within each configuration. (g) Rt density in the mBR relative to the mixed reactor. CC, co-culture; Ax, axenic control; mBR, membrane bioreactor; Se, *S. elongatus*; Rt, *R. toruloides*.

Growth and sucrose dynamics were mostly consistent with previous reports on Se– Rt co-culture (28). *S. elongatus* reached about 1.4 × 10^9^ cells mL^-1^ in all cultures by day 10 (**Fig. 1b, c**). Sucrose accumulated in both axenic controls, to 1.98 g L^-1^ in the mixed reactor and 1.77 g L^-1^ in the Se chamber by day 9, whereas in the mixed co-culture it fell below the assay limit from day 6. In the mBR, sucrose reached 1.07 g L^-1^ in the co-culture Se chamber and 1.40 g L^-1^ in the cell-free chamber, but did not exceed 0.23 g L^-1^ in the Rt chamber (**Fig. S3**), indicating that sucrose crossed the membrane and was consumed by the yeast. A difference in *S. elongatus* density between co-culture and its matched control appeared only when the two organisms shared a volume (**Fig. 1f**). In the mixed reactor, co- culture density was about 9% above its axenic control when averaged over days 3 to 8, largest on day 4 and declining to about 4% by day 10. In the mBR Se chamber the two were indistinguishable over the same interval. Effect sizes, confidence intervals and tests for all growth comparisons are given in **Table S1**. On days 1 and 2, before *R. toruloides* was added, co-culture and axenic cultures were indistinguishable in both configurations.

*R. toruloides* grew in both configurations but with different dynamics (**Fig. 1d, g**). Behind the membrane, its density was 17 to 31% lower than in the mixed reactor on days 4 to 6, equaled that of the mixed reactor at 165.6 h, and was 45% higher by day 10. The membrane therefore delayed yeast growth without limiting its final density. *S. elongatus* density was 5 to 10% higher in the mBR co-culture than in the mixed axenic culture.

### Symbols and colors

Points are replicates (pale fill) and group means (full fill); fill darkens with incubation time. Colors in (a) to (d) mark the culture (blue, CC; orange, Ax; purple, Rt in the mBR chamber). In (e) and (f), teal marks the partner effect in the mBR Se chamber; in (e), the open bar is the total and grey is the reactor-type part. Dotted lines mark the addition of Rt after the day 2 sampling (44.65 h) and, in (d) and (g), the time at which Rt density in the mBR crosses that in the mixed reactor. **Replication and statistics.** n = 3 per culture and day; error bars are one standard deviation (log scale for cell density, arithmetic for sucrose); shaded bands in (f) and (g) are Welch 95% confidence intervals. Panels use incubation hours because sampling intervals varied. **Supplementary data.** Sucrose in the Rt chamber and the blank chamber of the axenic mBR, **Fig. S3**; growth rates and statistics, **Table S1**.

### Only the mixed co-culture diverged from its axenic control for both exometabolome and transcriptome over time

Principal component analysis of the exometabolome and the transcriptome showed the same pattern over time **(Fig. 2**). In the mixed reactor, co-culture and axenic exometabolomes overlapped on days 1 and 2, separated after *R. toruloides* was added, separated further between days 5 and 6, and remained apart through day 9 (**Fig. 2a**). In the mBR, the co-culture Se chamber did not separate from the axenic Se chamber after day 3 (**Fig. 2b, c**), whereas the Rt chamber shifted again between days 6 and 7, close to the time at which yeast density behind the membrane exceeded that in the mixed reactor (**Fig. 1g**; **Fig. 2b**).

**Figure 2.**
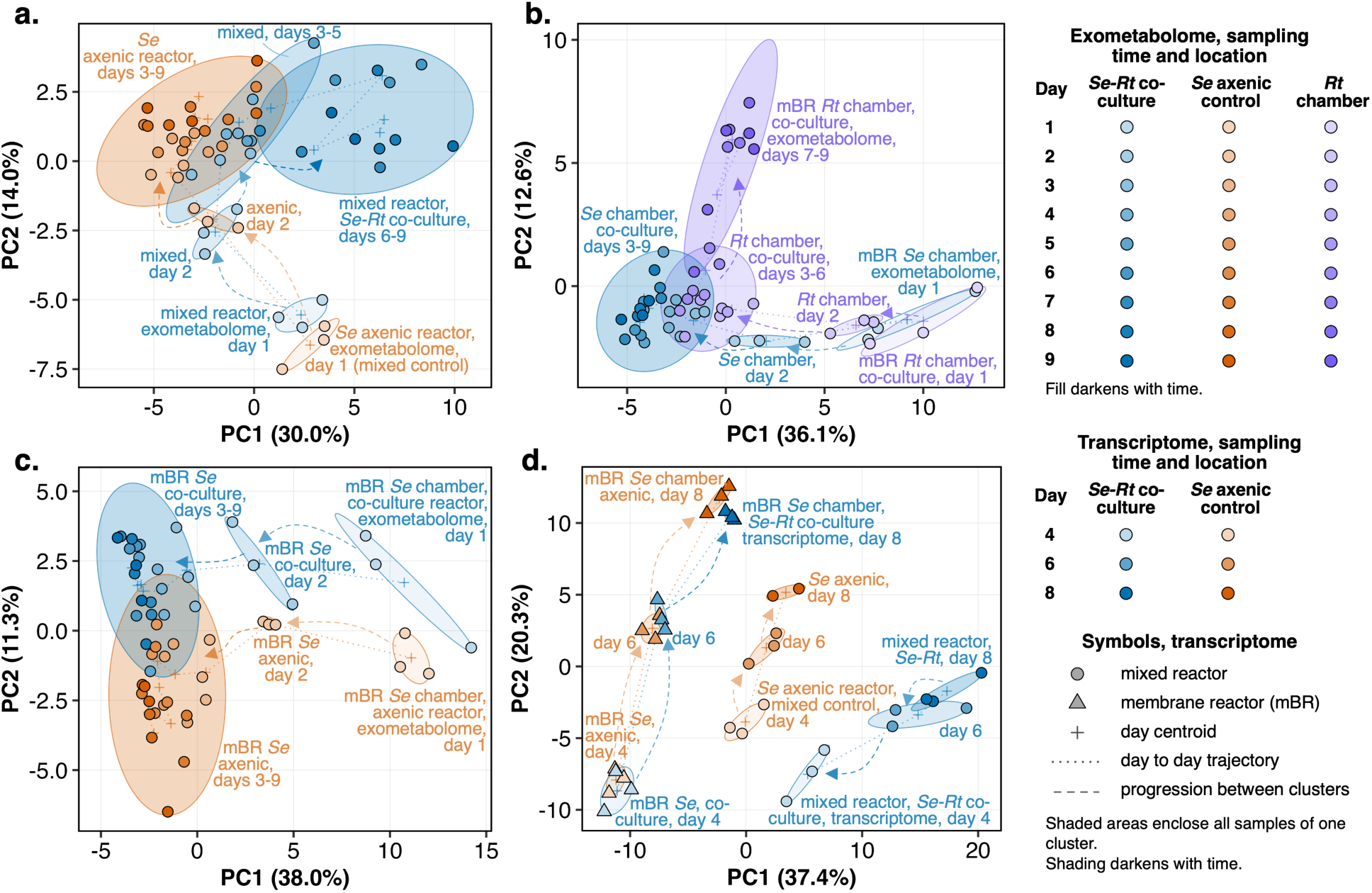
The exometabolome and the S. elongatus transcriptome diverged from the axenic control over time only in the mixed reactor. Principal component analysis of the exometabolome of (a) the mixed reactor, CC and Ax; (b) the CC mBR, Se and Rt chambers; (c) the mBR Se chamber, CC and Ax; and (d) the Se transcriptome of all four cultures on days 4, 6 and 8 (90.25, 138.25 and 188.25 h). CC, co-culture; Ax, axenic control; mBR, membrane bioreactor; Se, *S. elongatus*; Rt, *R. toruloides*. **Symbols and colors.** Symbols are individual samples, with fill darkening over time; crosses mark day centroids, dotted lines connect centroids, dashed arrows show progression between groups, and shaded areas enclose consecutive days that cluster together. Colors mark the culture as in Fig. 1: blue, CC; orange, Se Ax; purple, Rt chamber. Circles, mixed reactor; triangles, mBR. Rt was added after the day 2 sampling. **Replication and statistics.** Exometabolome data were log10(x + 1) transformed and Pareto scaled within the samples of each panel; the transcriptome was transformed with the DESeq2 variance-stabilizing transformation over all genes (35 samples).

The *S. elongatus* transcriptome followed the same pattern (**Fig. 2d**). The co-culture and axenic Se chamber cultures overlapped on all three sampling days, whereas the mixed co-culture separated from its control, and the separation increased from day 4 to day 8. The mixed axenic culture was also separated from both mBR cultures, consistent with an effect of the reactor configuration that is quantified below.

### Co-localization depleted the exometabolome, whereas membrane separation led to accumulation

The direction of exometabolome change differed between configurations (**Fig. 3a**). Across all 98 GC-MS metabolites, the mixed co-culture had lower levels than its axenic control (median log10 fold change -0.13, with 76% of compounds below zero; Wilcoxon signed-rank p = 4 × 10^-7^). Behind the membrane, the co-culture Se chamber had higher levels than its control (+0.13, p = 3 × 10^-8^), as did the Rt chamber relative to cell-free medium (+0.11, p = 0.006). The configuration control showed no overall shift (median -0.05, p = 0.58), although 68 individual metabolites differed between the two reactor types (**Fig. 3c**). Because the direction of change depends on the data treatment, it was determined from fixed-volume samples without sample normalization (**Methods; Text S1**).

**Figure 3.**
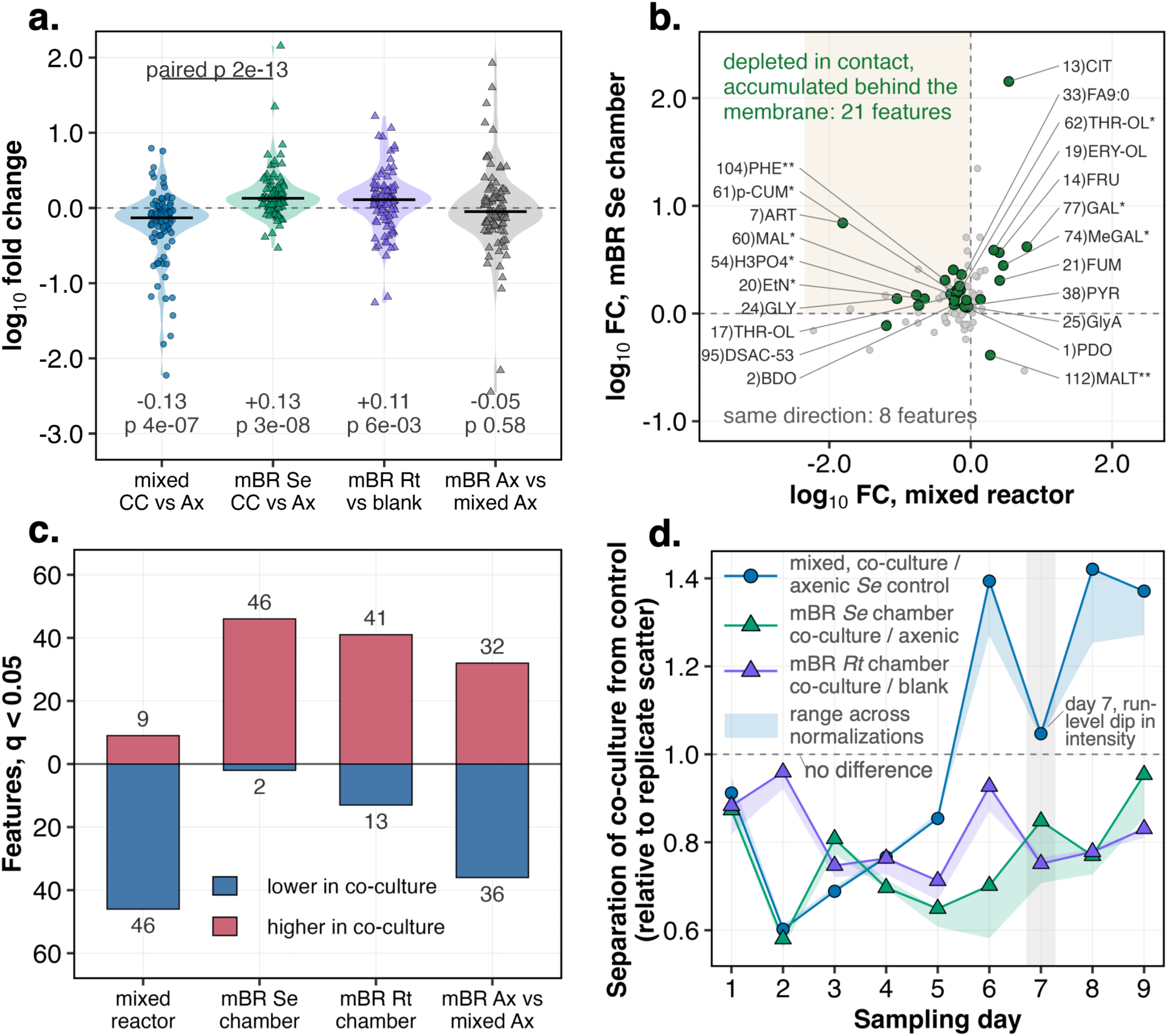
The exometabolome decreased with co-localization and increased behind the membrane. (a) Distribution of log10 fold change for all 98 compounds in each comparison; the bar is the median with a Wilcoxon signed-rank test against zero, and the bracket gives the paired test between mixed CC vs Ax and mBR Se CC vs Ax. (b) Compound by compound, mixed CC vs Ax against mBR Se CC vs Ax; filled symbols reach q < 0.05 in both (green), and shading marks depletion in direct contact with accumulation behind the membrane. Named compounds are labeled by code, and markers give identification confidence (**Table S2**). (c) Compounds reaching q < 0.05, lower (blue, below the axis) or higher (red, above) in the first group. (d) Distance between CC and Ax centroids relative to the mean distance between replicates (1, dashed line, means no separation); the band spans the three normalization treatments, and the shaded band at day 7 marks a run-level drop in total ion signal (samples retained, **Text S1**). Citrate (CIT) is higher in mixed CC; its large fold changes in the membrane comparisons reflect loss from the axenic Se chamber (**Text S1**). CC, co-culture; Ax, axenic control; mBR, membrane bioreactor; Se, *S. elongatus*; Rt, *R. toruloides*. **Symbols and colors.** Colors mark the comparison (first group minus second), not the culture as in Figs. 1 and 2: blue, mixed CC vs Ax; teal, mBR Se CC vs Ax; purple, mBR Rt vs blank (the Rt chamber against the cell-free chamber of the axenic mBR); grey, mBR Ax vs mixed Ax. **Replication and statistics.** Panels (a) to (c) use the primary analysis without sample normalization. **Supplementary data.** Non-detect counts for named compounds, **Table S2**.

Individual metabolites followed the same pattern (**Fig. 3b, c**). At q < 0.05 (Benjamini-Hochberg adjusted *p*), 46 compounds were lower and 9 higher in the mixed co- culture, mostly sugars and sugar alcohols together with the organic acids citrate and fumarate, whereas 2 were lower and 46 higher in the co-culture Se chamber. Of the 30 significant in both comparisons, 21 were depleted in the mixed reactor and accumulated behind the membrane. Metabolites depleted in the mixed reactor and detected in every sample included sucrose, two unidentified disaccharides (DSAC-52 and DSAC-50) and two phosphate compounds (H3PO4 and H3PO4*), consistent with sucrose consumption by the yeast and phosphate drawdown. Sugars were also drawn down behind the membrane: sucrose in the co-culture Se chamber was about 40% lower than in the axenic chamber by enzymatic assay (**Fig. 1c**), and the only two metabolites lower in this chamber were disaccharides (DSAC-53 and MALT**). The two configurations therefore differed mainly in the remaining metabolites, which decreased with co-localization and increased behind the membrane.

Exchange across the membrane was bidirectional. Of the 46 metabolites higher in the co-culture Se chamber, 32 were also higher in the Rt chamber than in cell-free medium, and 23 of these were detected in every mBR sample (**Table S2**), consistent with yeast- derived metabolites reaching *S. elongatus* across the membrane.

The exometabolome of the mixed co-culture diverged from its control progressively (**Fig. 3d**). The separation between co-culture and control, scaled to the scatter between replicates, remained between 0.6 and 0.9 over days 1 to 5, increased to 1.39 on day 6 and remained near 1.4 on days 8 and 9. Neither membrane comparison exceeded 0.96. Day 7 coincided with a run-level decrease in total ion signal, and excluding this day did not change any conclusion (**Text S1**).

### *S. elongatus* gene expression responded only in the mixed reactor

The *S. elongatus* transcriptome showed the same asymmetry as the exometabolome (**Fig. 4**). Because each reactor was sampled on all three days, reactor identity was fitted as a random effect (Methods). For the co-culture effect averaged over the three days, 51 genes were differentially expressed in the mixed reactor (35 higher and 16 lower in co-culture; q < 0.05 and absolute log₂ fold change above 1), compared with one gene in the mBR Se chamber (SYNPCC7942_RS13970, old locus tag Synpcc7942_0905, a predicted secreted protein of unknown function; +1.09). All 51 genes changed in the same direction on every day, and 42 reached q < 0.05 on each day. No gene was differentially expressed in both co- culture comparisons, and their fold changes were weakly correlated (r = 0.12; **Fig. 4b**). For the 55 genes with a significant configuration-by-culture interaction, the median absolute log2 fold change was 1.18 in the mixed reactor and 0.11 behind the membrane (**Fig. S4**), suggesting that the membrane response was close to absent rather than below the threshold. An independent analysis with DESeq2 recovered 48 of the 51 mixed-reactor genes (**Table S3**).

**Figure 4.**
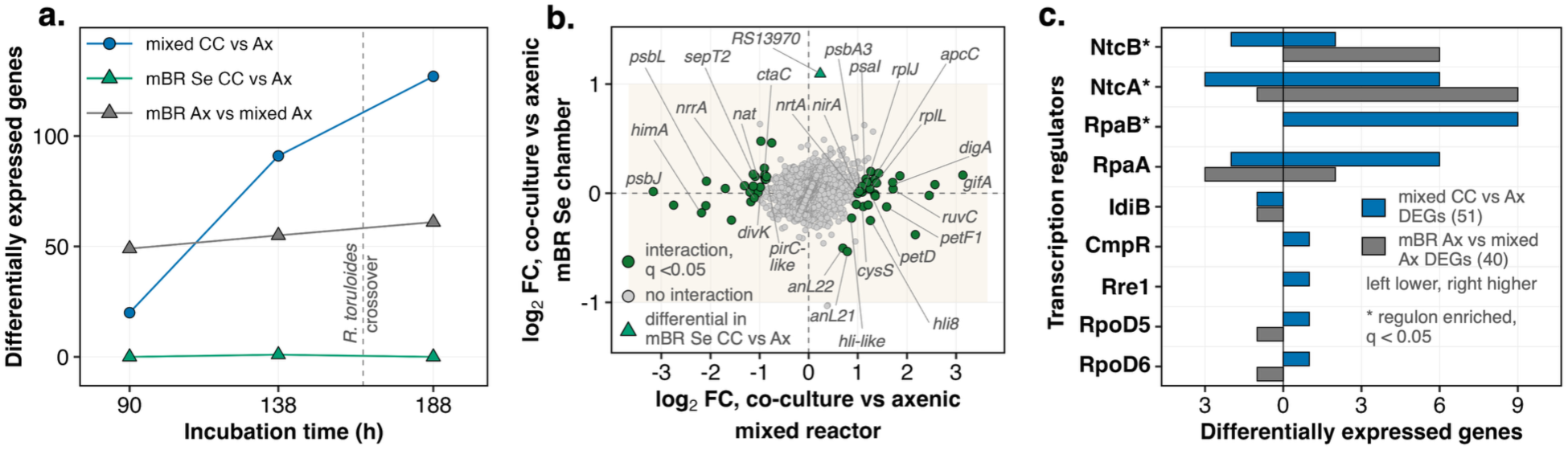
Co-culture altered S. elongatus gene expression in the mixed reactor but hardly at all behind the membrane. (a) Genes reaching q < 0.05 with an absolute log2 fold change above 1, tested within each sampling day. (b) Gene by gene, mixed CC vs Ax against mBR Se CC vs Ax, both averaged over days; filled symbols reach q < 0.05 for the configuration by culture interaction (green), the single gene differential in the mBR Se chamber is shown separately (teal triangle), and shading marks fold changes below the threshold in mBR Se. (c) Regulators of the differentially expressed genes in mixed CC vs Ax (blue) and mBR Ax vs mixed Ax (grey), from a curated network of 701 documented links; bars count genes higher (right) or lower (left) in the first group, and an asterisk marks regulons enriched in at least one set (one- sided Fisher test, q < 0.05). NtcB links and several NtcA links rest on binding-site evidence only. CC, co-culture; Ax, axenic control; mBR, membrane bioreactor; Se, *S. elongatus*; Rt, *R. toruloides*. **Symbols and colors.** Colors mark the comparison, not the culture as in Figs. 1 and 2: blue, mixed CC vs Ax; teal, mBR Se CC vs Ax; grey, mBR Ax vs mixed Ax. Genes are labeled by gene name where one exists, otherwise by RefSeq locus tag. **Replication and statistics.** The analysis covers 35 samples from days 4, 6 and 8 and 2,760 genes passing the expression filter. Reactor identity is a random effect, because each reactor was sampled on all three days; mixed Ax day 8 has n = 2 after one sample was excluded. The dashed line at 165.6 h in (a) marks where Rt density in the mBR equals that in the mixed reactor. **Supplementary data.** Day-averaged counts, sensitivity analyses, day-dependence tests and gene tables, **Table S3**; the 20 top interaction genes, **Fig. S4** and **Table S3**.

The response in the mixed reactor increased over the course of the experiment (**Fig. 4a**). Tested separately on each day with the same thresholds, 20, 91 and 127 genes were differentially expressed on days 4, 6 and 8, compared with 0, 1 and 0 in the Se chamber. Day 8 followed the point at which yeast density in the mBR reached that in the mixed reactor (**Fig. 1g**), so the difference between configurations is not explained by partner abundance. The configuration control changed 49 to 61 genes per day (40 averaged over days), a large and stable configuration effect that shared only 8 genes with the mixed- reactor response.

Regulator assignments indicated that the partner and the reactor configuration engaged different regulatory branches (**Fig. 4c**). Of the 51 mixed-reactor genes and the 40 configuration-control genes, 27 and 16 had a documented regulator. Both sets were enriched for NtcA targets (9 genes, q = 1 × 10^-4^; 10 genes, q = 2 × 10^-6^) and NtcB targets (4 genes, q = 1 × 10^-4^; 6 genes, q = 2 × 10^-8^). In the configuration control, the NtcB targets were the nitrate assimilation genes *nirA*, *nrtA* to *nrtD* and *narB*, all higher in the mBR. In the mixed reactor, NtcA targets changed in both directions, and *gifA*, which encodes the glutamine synthetase inactivating factor IF7, increased by 3.14 log2. RpaB targets were enriched only in the mixed reactor (9 genes, all higher; q = 0.005).

### Regulatory programs of *S. elongatus* changed only in the mixed reactor

Differential iModulon activity (DiMA), computed from the same samples with the *S. elongatus* PCC 7942 iModulon model (33), confirmed this asymmetry at the level of regulatory programs **(Fig. 5a)**. Nine iModulons differed between co-culture and axenic culture in the mixed reactor on at least two of the three days (raw p < 0.1): RpaB PS, RpaB∼P, PSII D1 and *psbJL* (photosynthesis and light response), NtcA IF7-17 (nitrogen), Fur-2 and CopR (metal homeostasis), cro and S-12. None differed in the Se chamber. *psbJL* decreased earliest and most strongly (-7.95, -9.61 and -10.92 on days 4, 6 and 8) and was the only iModulon that passed Benjamini-Hochberg correction (q = 0.009, 0.003 and 0.001). RpaB PS and PSII D1 increased, consistent with the opposite changes of *psbJ* and *psbL* (lower) and *psbA3*, *psaI* and *petD* (higher) at the gene level **(Fig. 4b)**. Fur-2 and S-12 also differed between the two axenic reactors and therefore reflect the reactor configuration as well as the partner.

**Figure 5.**
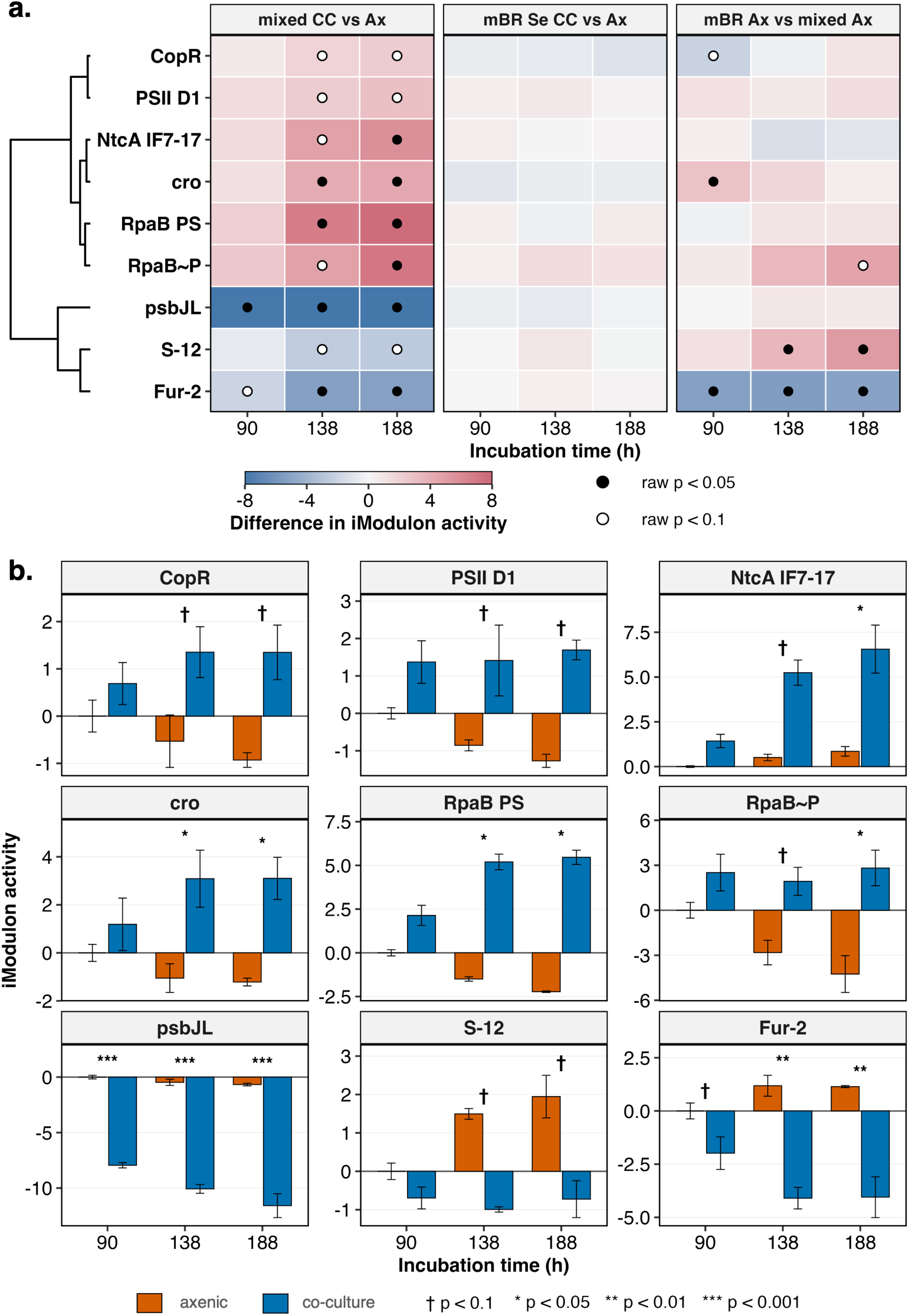
iModulon activity of S. elongatus changed with co-culture only in the mixed reactor. (a) Differential iModulon activity (first group minus second) for the nine iModulons that reach raw p < 0.1 on at least two of the three days in mixed CC vs Ax, by sampling day (90, 138 and 188 h); rows are ordered by Ward clustering of the mixed-reactor differences. (b) Activity in the mixed reactor, Ax and CC, by sampling day, relative to the mean of the three matched day-4 Ax flasks (zero by construction). CC, co-culture; Ax, axenic control; mBR, membrane bioreactor; Se, *S. elongatus*; Rt, *R. toruloides*. **Symbols and colors.** In (a), fill is the activity difference (blue, lower; red, higher in the first group), filled dots mark raw p < 0.05 and open dots raw p < 0.1. In (b), bars are group means, blue marks CC and orange marks Ax, as in Figs. 1 and 2; marks give the DiMA test (* p < 0.05, ** p < 0.01, *** p < 0.001, † p < 0.1). **Replication and statistics.** Error bars are one standard deviation. The analysis covers the 35 samples of Fig. 4; mixed Ax day 8 has n = 2 after one sample was excluded. Significance is shown on raw p values; after Benjamini-Hochberg correction only *psbJL* remains significant. **Supplementary data.** Activities in the mBR Se chamber, NtcA programs against two nitrogen-starvation references, and the perturbation table, **Figs. S5** an**d S6** and **Table S4**.

The nitrogen programs of the mixed co-culture moved away from a starvation pattern (**Fig. 5b**). NtcA IF7-17 increased in the mixed co-culture by +1.43, +4.74 and +5.71 relative to its control (raw p = 0.25, 0.061 and 0.047), while the axenic culture remained near its day-4 level. Two nitrogen-starvation references analyzed in the same way, wild- type *S. elongatus* and a related co-culture (**Fig. S5**), both decreased NtcA IF7-17 (-7.92 and -3.89) and increased NtcA-1 and NtcA-2. The mixed co-culture changed all three in the opposite direction, although only NtcA IF7-17 reached p < 0.05. The reactor configuration affected a different branch: NtcAB NO3, the nitrate assimilation iModulon, was higher in the mBR axenic culture (+3.76, +5.14 and +5.11; p = 0.062, 0.036 and 0.037), whereas NtcA IF7-17 did not differ between reactor types. iModulon activities in the Se chamber and all DiMA values are provided in **Fig. S6** and **Table S4**.

## Discussion

Metabolite exchange alone did not reproduce the response of *S. elongatus* to its partner. Sucrose and yeast-derived metabolites crossed the membrane, and *R. toruloides* reached a higher final density behind it, consistent with the shared-pool view for the yeast side of the interaction (14, 21). *S. elongatus*, however, changed its gene expression and regulatory programs only when the partners shared a volume. The response of a phototroph to a heterotrophic partner varies between systems (30); here it depended on co-localization. Two explanations are consistent with these data: *S. elongatus* may respond to a signal that does not enter the diffusible pool, or a diffusible signal may be diluted below an effective concentration before it crosses the membrane. Candidate mechanisms include steep chemical gradients near the cell surface (2), attachment and joint extracellular matrices (15, 22), and competition for shared resources within one volume. The present design does not distinguish among these mechanisms; co-cultures that permit contact while limiting bulk exchange, or co-culture with inactivated partner cells, would allow this distinction.

The effect of membrane separation is likely to depend on what the partner supplies and what limits the phototroph. Across a membrane, heterotrophic bacteria grown on 5 g L⁻¹ glucose without CO2 supply have been reported to increase *S. elongatus* density several-fold, with a large accompanying change in its transcriptome (29). Under those conditions, respiratory CO2 and bacterial by-products derived from the added glucose can cross the membrane and may relieve carbon limitation of the phototroph. In the present system, the reactors were supplied with 2% CO2, and *R. toruloides* depended on sucrose produced by *S. elongatus*, so this route was largely absent. Together, the two studies suggest that diffusible exchange can account for the phototroph response when it delivers a limiting resource, whereas without such a resource the response requires co-localization.

A growth benefit attributed to a partner can instead arise from differences in reactor configuration. Here a partner effect on *S. elongatus* density was detected only in the mixed reactor (**Fig. 1f**), whereas the axenic mBR culture was already denser than the mixed axenic culture (**Fig. 1e**). Separating the two effects also accounts for the growth pattern of our earlier study, in which *S. elongatus* grew better in membrane-separated than in mixed co- culture (28). The difference may reflect the timing of yeast addition: at inoculation in the earlier study, and here 44.65 h after induction, when *S. elongatus* had reached 6.8 × 10^8^ cells mL⁻¹. Phototroph-heterotroph interactions can shift from cooperation to competition with nutrient availability and growth stage (19, 21). A yeast added at inoculation, before sucrose secretion is established, has nearly no carbon source, and this timing may change how it interacts with its partner. Varying the time of yeast addition would test whether timing determines the outcome.

The opposite directions of exometabolome change are consistent with a balance between release and removal. In a shared volume, the yeast has immediate access to released metabolites, so uptake keeps pace with release and extracellular concentrations decrease. Behind the membrane, transport limits removal, and metabolites accumulate on the side where they are produced; sucrose follows this pattern (**Fig. 1b, c**). Accumulation in the Rt chamber relative to cell-free medium likely also includes metabolites released by the yeast. The three readouts followed different time courses: the growth difference in the mixed reactor peaked early and faded, whereas exometabolome separation and the number of differentially expressed genes increased to day 8. The exometabolome integrates release and uptake up to each sampling point, and the transcriptome followed this slower course. The time dependence is consistent with observations in *Prochlorococcus-Alteromonas* co- cultures (19, 20) and suggests that a single sampling point could miss the difference between configurations or a change in the nature of the interaction over time.

The regulatory programs point to nitrogen status and photosynthesis as the main targets of the response to co-localization. NtcA integrates carbon and nitrogen status (34). In the mixed co-culture, NtcA IF7-17, whose leading genes *gifA* and *gifB* encode inactivating factors of glutamine synthetase (35), increased, whereas NtcA-1 and NtcA-2 decreased, opposite to two nitrogen-starvation references. The *pirC*-like gene was also lower by 0.87 log₂; its *Synechocystis* homolog has been reported to be activated by NtcA and strongly induced under nitrogen starvation (36, 37). Together, these patterns are consistent with *S. elongatus* in the mixed co-culture not being nitrogen-limited. Whether this reflects nitrogen released by the yeast, a change in carbon status, or both cannot be resolved, because nitrogen species were not measured. The reactor configuration affected a separate branch, nitrate assimilation, distinguishing the two nitrogen responses. Among photosynthesis programs, *psbJL* decreased on all three days, while RpaB PS, PSII D1 and several photosystem genes increased, suggesting remodeling of the photosynthetic apparatus rather than a uniform change in its expression. Light availability in a culture shared with yeast cells was not measured here.

These results are relevant to the design of phototroph-heterotroph consortia for bioproduction (12, 13). Membrane separation allows each partner to be harvested and controlled independently and prevents one partner from overgrowing the other. Where the aim is one-way carbon delivery to the heterotroph, separation had little cost here, as the yeast reached a 45% higher density behind the membrane by day 10. Where the aim depends on benefits returned to the phototroph, such as improved growth, stress protection or viability reported in mixed co-cultures (11, 16, 17), co-localization may be required unless the heterotroph supplies a diffusible limiting resource.

The design used here separates the partner effect from that of the reactor configuration and follows the response over time at four levels, so the main result, a response of *S. elongatus* confined to co-localization, is supported by consistent growth, exometabolome, gene expression and regulatory program data, and the approach is applicable to other membrane-separated co-cultures. Several aspects define the scope of the conclusions. The mixed reactor shares nitrogen, phosphorus, inorganic carbon and a single light environment in addition to permitting cell contact, so the comparison estimates the combined contribution of all factors excluded by the membrane rather than the effect of contact alone. The transcriptome analysis covers *S. elongatus* only, so the regulatory response of the yeast and the coordination between the cyanobacterial and yeast transcriptomes remain to be determined. Exchange across the membrane was not quantified as fluxes. Thirty-eight of the 98 exometabolome compounds are unidentified, which limits chemical interpretation but not the direction of change. Finally, the results come from one cultivation experiment on one phototroph-heterotroph pair; a replicate experiment and other partnerships would test the generality of the findings.

## Materials and Methods

### Strains, medium and cultivation

The sucrose-secreting *S. elongatus* PCC 7942 *cscB*/SPS strain, which expresses the sucrose permease CscB and sucrose phosphate synthase under IPTG control (38), was provided by D. Ducat (Michigan State University). *R. toruloides* NBRC 0880 (IFO 0880) was the yeast partner. Medium, temperature, CO2 supply, stirring and IPTG induction were as described (28): BG11 supplemented with yeast nitrogen base without amino acids and ammonium sulfate (0.09 g L⁻¹), ammonium sulfate (0.264 g L⁻¹), K2HPO4 (0.174 g L⁻¹) and HEPES (3 g L⁻¹), pH 8.4; 30 °C; 2% CO2 in N2; magnetic stirring at about 200 rpm; continuous light at about 200 µmol photons m^-2^ s^-1^ from 15 W LED panels. *S. elongatus* inoculum cultures were grown in the same medium with 25 mg L^-1^ kanamycin and 12.5 mg L^-1^ chloramphenicol. Cells were harvested by centrifugation, washed and resuspended in antibiotic-free medium with 1 mM IPTG. *R. toruloides* precultures were grown in the same medium with 10 g L^-1^ sucrose, then harvested and washed.

### Reactor configurations and sampling

Two-chamber mBRs were built as described (28), from modified 500 mL and 250 mL GL45 glass bottles joined at NW63 flanges holding a 0.22 µm hydrophilic PVDF membrane (Durapore, Millipore). The *S. elongatus* chamber held 665 mL of culture and the *R. toruloides* chamber 425 mL (**Fig. S1 and S2**). The mixed reactor was a single borosilicate glass bottle with working volume about 500 mL. Four cultures were run with three reactors each: mixed co-culture, mixed axenic, mBR co-culture (Se chamber and Rt chamber) and axenic mBR (Se chamber and a second chamber holding cell-free medium). The two configurations were inoculated with *S. elongatus* at 47.9 × 10^6^ (mixed) and 37.0 × 10⁶ (mBR) cells mL⁻¹. *R. toruloides* was added to the mixed co-culture and to the Rt chamber at 44.65 h, after the day 2 sampling, to about 5 × 10⁶ cells mL⁻¹. Each day for 10 days, 1 mL was withdrawn for cell counts and 3 mL for exometabolome and transcriptome analysis, or 6 mL from the *R. toruloides* chamber. Biomass was collected by centrifugation, the supernatant was passed through a 0.2 µm filter, and both fractions were frozen in liquid nitrogen.

### Cell counts and sucrose

*S. elongatus* and *R. toruloides* cells were counted by flow cytometry (NovoCyte 3000 VYB, Agilent), distinguished by forward scatter and the red autofluorescence (660/20 nm) of photosynthetic pigments. Samples were run at 50 µL min⁻¹ to at least 100,000 *S. elongatus* events, with medium and autoclaved water as negative controls, and analyzed in NovoExpress (Agilent). For sucrose content, the filtered supernatant was assayed enzymatically (Sucrose/D-Glucose Assay Kit K-SUCGL, Megazyme).

### Exometabolome

Filtered supernatant samples were analyzed at the Environmental Molecular Sciences Laboratory (EMSL Award 60520). Samples were dried, derivatized by methoxyamination and trimethylsilylation, and analyzed by GC-MS on an Agilent 8890 GC system with a 5977 Inert Plus single-quadrupole MSD and an HP-5MS column (30 m × 0.25 mm × 0.25 µm, Agilent), with splitless injection at 250 °C and an oven ramp from 60 to 325 °C. Compounds were deconvoluted, aligned and identified against an augmented in-house library and the NIST23 and Wiley11 libraries with MetaboliteDetector (39). Identification confidence is carried in the compound codes used throughout: no symbol for a match score above 0.75, one asterisk below 0.75, and two asterisks for a best spectral match only. After removal of column bleed, QC failures and carryover, 98 compounds in 162 samples were analyzed (**Text S1; Table S2**).

Exometabolome data were not normalized per sample, because aliquots were of fixed volume; probabilistic quotient normalization (PQN) was run as a sensitivity analysis (Text S1). Differential abundance was tested for each compound with a linear model, log10(x + 1) ∼ group + day, across days 1 to 9, with Benjamini-Hochberg correction (^40^) across the 98 metabolites (q). Pool-level shifts used Wilcoxon signed-rank tests of the 98 fold changes against zero. Separation per day (**Fig. 3d**) is the distance between group centroids divided by the mean within-group distance, pooled across days, after log10(x + 1) transformation and Pareto scaling, and was tested by permuting labels within days (4,999 permutations). Principal component analysis (**Fig. 2a** to **c**) used the same transformation within each panel’s subset.

### Growth statistics

Co-culture effects on *S. elongatus* density are geometric mean ratios of co-culture to axenic density on the same day, with Welch 95% confidence intervals on ln density. For the pooled estimate over days 3 to 8, each reactor was averaged over those days before the cultures were compared (n = 3 reactors per culture). Growth rates and curve fits are described in **Text S1.**

### RNA sequencing and differential expression

*S. elongatus* cells were harvested at 90.25, 138.25 and 188.25 h as above. RNA was extracted with the RNeasy kit and QIAshredder columns (QIAGEN) after lysis in RLT buffer with 1% β-mercaptoethanol, treated with DNase and depleted of rRNA. Paired-end reads (2 x 150 bp) were sequenced on an Illumina platform by Azenta US, Inc., aligned with Rsubread to the PCC 7942 RefSeq genome and counted as described (33). Samples carried 38 to 89 million gene-assigned reads. One mixed axenic sample from day 8 departed from its two replicates (Spearman r 0.983 and 0.980, against 0.994 between those two) and was excluded, leaving 35 samples (**Text S1**).

Genes with at least 10 counts in at least 3 samples were kept (2,760 of 2,761) and normalized by the trimmed mean of M values (41). Differential expression used limma- voom (42) with the twelve culture-by-day groups as a means model. Because each reactor was sampled on all three days, reactor identity was fitted as a random effect with duplicateCorrelation (43) (consensus correlation 0.124). Day-averaged and per-day contrasts came from the same fit, and a gene was called differentially expressed at q < 0.05 and absolute log2 fold change above 1. Day dependence of the co-culture effect was tested by a joint F test on the day 6 minus day 4 and day 8 minus day 4 differences. The configuration by culture interaction is the mixed-reactor contrast minus the Se chamber contrast. DESeq2 (design ∼ day + culture, Wald test) served as an agreement check (**Table S3**). The transcriptome PCA (**Fig. 2d**) used the DESeq2 variance-stabilizing transformation of all genes.

Enrichment used one-sided Fisher exact tests against all expressed genes, with Benjamini-Hochberg correction within each family (regulons, curated *S. elongatus* gene sets, KEGG pathways and COG categories; set sizes and thresholds in **Text S1**). Regulons were defined using the 701-link transcriptional regulatory network of the *S. elongatus* PCC 7942 iModulon model (33). Proteins of differentially expressed genes without an assigned function were re-annotated from domain, structure and topology predictions (**Text S1**).

## iModulon activities

Each sample was projected onto the 78 iModulons of the *S. elongatus* PCC 7942 model, built from 407 RNA-seq samples with optICA (44). Differential iModulon activity (DiMA) was computed for each day and comparison against a log-normal null fitted to replicate differences within the 177 distinct experimental conditions of the compendium (32). The day-4 axenic control was the three matched mixed axenic samples of this experiment, and the excluded day-8 sample was left out. iModulons were shown in **Fig. 5** if raw p was below 0.1 on at least two of the three days in the mixed reactor; Benjamini-Hochberg adjusted p is reported alongside. Absolute activities (**Fig. 5b**) are relative to the mean of the matched day-4 axenic samples. Two nitrogen-starvation references were computed with the same null: wild-type *S. elongatus* after 48 h without nitrogen against nitrogen-replete wild type (SRR3283837 to SRR3283840), and a *S. elongatus* and *R. toruloides* co-culture after 2 h without nitrogen against the same co-culture with nitrogen (**Table S4**).

## Software

Transcriptome analyses and figures used R 4.6.0 with limma 3.68.5, edgeR 4.10.3 (45), statmod 1.5.2, DESeq2 1.52.0 (46) and ggplot2 4.0.3, and Rsubread 2.20 (47). Exometabolome, growth and DiMA calculations used Python 3.12.15 with NumPy 2.5.4, pandas 3.0.6 and SciPy 1.18.1. DiMA followed the PyModulon approach (32).

## Data, Metadata, and Code Availability

RNA-seq raw sequencing data are publicly available under the GEO accession GSE335065 and under NCBI BioProject accession PRJNA1477361. GC-MS raw data and the processed metabolite table are in MassIVE (MSV000099916). Cell counts, sucrose, the exometabolome matrix, all result tables and sample metadata following STREAMS are provided in **Tables S1** t**o S4** and in the Zenodo archive. All analysis and figure scripts are available at https://github.com/PBohutskyi/cyanobacteria-yeast-coculture-mixed-vs-mBR, archived with a DOI at Zenodo (doi:10.5281/zenodo.23456462).

## Acknowledgments

This work was supported by the Predictive Phenomics Initiative under the Laboratory Directed Research and Development Program at Pacific Northwest National Laboratory (PNNL), and in part by the BRaVE project funded by the U.S. Department of Energy (DOE), Office of Science, Office of Biological and Environmental Research (BER), under FWP 81832. A portion of this research was performed under the DOE Office of Science Reaching a New Energy Sciences Workforce (RENEW) contract DE-SC0024289. A portion of this research was performed under EMSL Award 60520 (https://doi.org/10.46936/cpcy.proj.2022.60520/60008548) at the Environmental Molecular Sciences Laboratory, a DOE Office of Science User Facility sponsored by BER.

Z.J. was supported by the PNNL–Washington State University Distinguished Graduate Research Program (DGRP) Fellowship. We thank Daniel C. Ducat (Michigan State University) for providing the *S. elongatus* PCC 7942 cscB/SPS strain. PNNL is operated by Battelle for the DOE under Contract DE-AC05-76RL01830.

## Author Contributions

**Pavlo Bohutskyi:** Conceptualization, Methodology, Investigation, Formal analysis, Visualization, Data curation, Funding acquisition, Project administration, Writing – original draft, Writing – review & editing, Supervision. **Zachary Johnson:** Formal analysis (transcriptomics, iModulon analysis), Software, Writing – review & editing. **Marija Veličković:** Investigation (metabolomics), Writing – review & editing. **Young-Mo Kim:** Investigation, Formal analysis (GC-MS metabolomics), Writing – review & editing. **Margaret S. Cheung:** Funding acquisition, Supervision, Project administration, Writing – review & editing.

