## Supplementary material for "Metabolites cross the membrane, the transcriptome does not follow: co-localization is required for the cyanobacterial response to a yeast partner": Text S1

Supplementary Text S1, Figures S1 to S6 and legends for Tables S1 to S4

*Companion files: Table_S1_growth.xlsx, Table_S2_exometabolome.xlsx, Table_S3_transcriptome.xlsx, Table_S4_iModulons.xlsx*

### Contents

Contents: Text S1 (supplementary methods and notes); **Figures S1 to S6** with legends; and the legends for **Tables S1 to S4**. Compact summary tables are reproduced below as Word tables; the complete data for every **Table Sx** is provided in the matching Excel workbook (Table_Sx.xlsx), one named sheet per block of data.

#### Text S1

##### *Cultivation, sampling and time base*

Four cultures, three biological replicates each: mixed co-culture, mixed axenic *S. elongatus*, two-chamber membrane bioreactor (mBR) co-culture, and two-chamber mBR with axenic *S. elongatus* and a blank second chamber holding medium only. The two-chamber configuration matches the one used in Bohutskyi et al. 2024 (1); the membrane-format axenic control takes the place of the three-chamber configuration of that study.

Cell counts were collected at 11 time points (days 0 to 10) and exometabolome samples at 9 (days 1 to 9). Sampling intervals were 21.6 to 25.25 h, so all rates, fits and figures use elapsed time; day labels are kept in sample names and text (**Table S1**, sheet Time_base).

##### *Growth*

Specific growth rates were estimated two ways (**Table S1**): point to point on group geometric means over each interval (method A), and by log-linear regression over a fixed window (method B; days 1 to 3 for *S. elongatus*, days 3 to 6 for *R. toruloides*). The membrane-format cultures have a clean exponential window ($R^2 > 0.99$). In the mixed format growth is nearly complete by day 3, no window reaches $R^2$ above 0.88, and the estimate depends on a single interval, so the mixed rate is reported but not compared with the membrane rate as a point estimate.

Co-culture effects are geometric mean ratios of co-culture to axenic density on the same day, with Welch 95% confidence intervals on ln density, n = 3 per group, BH across days within each configuration. Because the three replicates are three reactors sampled repeatedly, the pooled estimate over days 3 to 8 is computed at reactor level: each reactor is averaged over those days, then reactors are compared, n = 3 per group (**Table S1**, sheet Pooled_days3to8).

The two configurations received different inocula, 47.9 and 37.0 × $10^6$ cells per mL, so every co-culture comparison is made within one configuration, and the raw density ratio between configurations (sheet Se_mBRAx_vs_mixedAx) carries the inoculum difference. *R. toruloides* has no axenic control by design, so for the yeast a configuration effect cannot be separated from a partner effect; densities are per mL of the chamber the organism occupies. Curve fits (Zwietering-form logistic and Gompertz on ln(N/N0)) use the day-2 count as N0 for *R. toruloides*, which was not counted on days 0 and 1; a three-parameter logistic on absolute density is given alongside.

##### *Exometabolome processing*

GC-MS of filtered spent medium, 2 mL fixed-volume aliquots, methoxyamination and trimethylsilylation, single quadrupole detection. The analysis matrix is 98 compounds by 162 samples (6 groups x 9 days × 3 replicates). Non-detects are written as explicit zeros and number 488 (3.07% of the matrix); 63 metabolites are detected in every sample and none falls below 50% detection. Fourteen of the 112 compounds in the raw table were removed: 1 chemical (column bleed), 2 by the QC screening rule, and 11 as injection carryover from plant samples run immediately before this batch. Eleven compounds were renamed where the assigned name could not be correct. Identification markers and the source of each retention time, signal-to-noise ratio and library score are given in **Table S2** (sheet Features and README). The marker convention differs from the one used in Bohutskyi et al. 2024 and should be read from **Table S2**.

Sucrose. 42)SUCR carries 86.7% of the total ion signal at the median sample and up to 94.1%. Its response compresses at the top of the range, giving a mixed axenic over co-culture ratio of about 18-fold pooled and 31-fold at day 9, against more than 100-fold measured by enzymatic assay. Sucrose magnitudes are therefore taken from the enzymatic assay (**Figure 1**), with the GC-MS peak cited for identification.

Day-7 intensity dip. Total ion signal excluding 42)SUCR and 95)DSAC-53 (a disaccharide that tracks sucrose) falls at day 7 in 5 of the 6 groups and recovers at day 8 in 5 of 6 (**Table S2**, sheet Total_ion_signal). Pooled across groups, day 7 sits at 81% of the mean of days 6 and 8, and the dip is present in all three replicate blocks. The replicate blocks were injected consecutively, so day 7 holds the seventh injection position in every block; the pattern is consistent with a run-level effect rather than a biological event. The dip lowers the between-group distance and raises the within-group scatter on that day, and normalization does not remove it. Day 7 is retained in every statistic: the differential abundance model blocks on day, and the permutation test shuffles labels within each day. In **Figure 3d**, day 7 is marked by a shaded band; the point is drawn like the others. Excluding day 7 changes no conclusion: mean divergence over the run moves from 1.01 to 1.00 in the mixed reactor, 0.76 to 0.75 in the mBR Se chamber and 0.82 to 0.83 in the mBR Rt chamber.

##### ***Exometabolome statistics***

Each analysis subsets to the samples in the comparison before transforming, so variance from cultures outside the comparison never enters the scaling. Divergence uses $\log_{10}(x + 1)$ followed by Pareto scaling within the subset; differential abundance uses $\log_{10}(x + 1)$ only. A compound-wise linear model is invariant to Pareto scaling (largest difference in t across the 98 metabolites $2 \times 10^{-13}$), so fold changes stay in $\log_{10}$ units.

Divergence per day is the distance between group centroids divided by the mean within-group distance, both in the Pareto-scaled space, with the denominator pooled across days so that the metric does not move when one day happens to be unusually tight or noisy (**Table S2**, sheet Divergence). With n = 3 against 3 there are only 10 distinct label splits per day, so per-day divergence is reported as an effect size, and significance is tested once per comparison by permuting labels within every day simultaneously (4,999 permutations). Differential abundance uses $\log_{10}(x + 1) \sim \text{group} + \text{day}$ across all nine days, with the t test on the group term and BH correction across the 98 metabolites (**Table S2**, sheet Fold_changes).

Normalization. The primary analysis uses no sample normalization: aliquots were of fixed volume, technical scale variation is 13% CV within replicate cells, and the main assumption of probabilistic quotient normalization (PQN), that most metabolites are unchanged, does not hold in the mixed reactor, where 74 of 98 compounds are lower in the co-culture. PQN against a stable 20-compound reference is the designated sensitivity analysis, and PQN over all 98 compounds an additional check (**Table S2**, sheet Normalization). Full PQN converts the 46 to 9 depletion in the mixed reactor into a 24 to 25 split, and the 2 to 46 accumulation in the mBR Se chamber into 20 to 18. The mBR Rt chamber, where 32% rather than 76% of compounds are lower, is less affected. The divergence magnitude is robust to the choice; the direction is not.

Non-detects. Among compounds at $q < 0.05$, non-detects occur in 10 of 55 (mixed reactor), 9 of 48 (mBR Se chamber), 17 of 54 (mBR Rt chamber) and 16 of 68 (mBR Ax vs mixed Ax); at least 5 in one group in 5, 3, 8 and 7 (**Table S2**, sheets Nondetects_named and Nondetects_all). In the mBR Rt chamber comparison the control chamber holds cell-free medium, so a compound not detected there but detected with the yeast is a detection result rather than a graded fold change. In the Rt chamber, 7 of the 41 metabolites higher than cell-free medium were absent from 5 to 13 of the 27 cell-free samples. Restricted to compounds detected in every sample of a comparison (72 to 79 compounds), the median $\log_{10}$ fold change keeps its sign: mixed reactor -0.10 (Wilcoxon p = $8 \times 10^{-6}$), mBR Se chamber +0.10 ($p = 4 \times 10^{-6}$), mBR Rt chamber +0.09 ($p = 0.03$), mBR Ax vs mixed Ax -0.03 ($p = 0.85$).

Citrate. 13)CIT (library match, score 0.97) is detected in every group on days 1 and 2 and falls in all groups from day 3 (**Table S2**, sheet Citrate_by_day). In the axenic mBR Se chamber it falls to near the detection limit: from day 3, 13 of 21 samples are non-detects and 7 of the 8 detected values are below 5,000. The co-culture mBR Se chamber stays at levels similar to the mixed axenic culture (medians over the run 25,905 and 27,517), and the mixed co-culture holds the most (128,138). Both Rt-side chambers are mostly non-detects from day 7. The large mBR Se chamber (+2.15) and mBR Ax vs mixed Ax (-2.45) fold changes for citrate therefore reflect its loss in the axenic mBR Se chamber rather than a difference between co-culture and axenic culture. In the mixed reactor, where citrate was detected in every sample, it was higher in co-culture (+0.54, $q < 0.001$).

Compound notes.

- 15)GLC is reported as glucose following the manual check and assignment; the library call for that peak is D-galactose. Raw GC-MS confirms a clean, baseline-resolved peak carrying the co-culture decrease, and the library's basis for the galactose call, a major and minor oxime pair with 77)GAL, does not hold. The isomer remains unresolved: the spectrum is a generic aldohexose methoxime-TMS and no retention index ladder was available.
- 95)DSAC-53 (Unknown 053 in the EMSL matrix) elutes next to sucrose, 42)SUCR (library score 1.00, S/N 1471), and follows its trajectory closely, but its spectrum differs from sucrose. It is reported as an unidentified disaccharide.
- 103)3HPA** is a chromatographic component in its own right, not a split of a neighbouring peak, but the 3-hydroxypropanoic acid assignment is not supported by the diagnostic ions.
- 10)HCO3 is retained for its contribution to group separation and is not interpreted mechanistically, since the signal combines consumption with $CO_2$ sparging, added bicarbonate, buffering and extraction pH.
- 21)FUM and 70)FAL* carry a partial replicate-order gradient; no claim depends on either alone.
- 31)MALT* has no direct library match; maltose is a curated candidate only.

***Transcriptome: sensitivity and agreement checks***

Repeated sampling. Because reactors were sampled repeatedly, a culture holds three experimental units rather than nine. Two treatments of this structure were compared with a third in which it is ignored (**Table S3**, sheet Sensitivity). Ignoring it gave 52 differentially expressed genes in the mixed reactor; fitting reactor as a random effect, the analysis used in the paper, gave 51; summing each reactor across its three days before a single between-culture test, the analogue of the reactor-level growth analysis, gave 47. The mBR Se chamber gave 1 gene under all three.

Normalization. TMM was the primary normalization. Upper-quartile normalization, chosen because the mixed co-culture samples have the fewest reads, gave 52 genes in the mixed reactor against 51, and the two sets of fold changes correlate at $r > 0.999$.

Agreement with DESeq2. The four day-averaged comparisons were repeated with DESeq2 (design ~ day + culture, Wald test, adjusted $p < 0.05$ and $|\log_2 FC| > 1$), which does not model the repeated sampling of reactors. Fold changes agreed with limma at $r = 0.995$ to 0.998, and 48 of the 51 mixed-reactor genes, the single mBR Se chamber gene and 39 of the 40 configuration-only genes were called by both. With adaptive shrinkage of the DESeq2 fold changes (ashr), every DESeq2 gene is also a limma gene (**Table S3**, sheet limma_vs_DESeq2).

Excluded sample. Sample Se_ax_190_1 (mixed axenic, day 8) departed from both other replicates (Spearman r 0.983 and 0.980, against 0.994 between those two) and lay furthest from its group centroid of all 36 samples, while Cook's distance flagged no gene in any sample (**Table S3**, sheets Excluded_sample and Excluded_sample_pairs).

Enrichment. Tests covered 12 regulons with at least 6 genes, from a curated network of 701 documented regulator-target links; curated *S. elongatus* gene sets; 78 KEGG pathways with at least 8 genes; and COG categories with at least 10 genes.

***Re-annotation of genes without an assigned function***

Of the *S. elongatus* genes differentially expressed in any comparison, 31 had no assigned function in the RefSeq annotation (GCF_000012525.1). Their proteins were re-annotated with sequence- and structure-based searches. Domains and families were identified with InterProScan 5 (2) against all InterPro member databases (3) through the EMBL-EBI Job Dispatcher (4), and transmembrane helices and signal peptides were predicted with Phobius (5). Structural sequences were predicted with ProstT5 (6) and searched with Foldseek (7) against the PDB and AlphaFold/Swiss-Prot databases. A function was assigned from an informative domain name, or from the best informative structure match with an E-value of at most $10^{-5}$ covering at least half of the query. Proteins with neither were classed by predicted topology or as unknown (**Table S3**, sheet Reannotation).

Three proteins received a domain-based function, eight were predicted membrane or secreted proteins from topology alone, and 20 remained unknown. Among the 51 genes that changed in the mixed reactor, 18 lacked a function; one received a domain (RmlC-like cupin fold, SYNPCC7942_RS12290), four a predicted membrane or secreted location, and 13 remained unknown. The single gene that changed in the mBR Se chamber, SYNPCC7942_RS13970, carries a predicted signal peptide. SYNPCC7942_RS05895, a DUF305 protein with a predicted signal peptide that was higher in the mixed co-culture on days 6 and 8 ($log_2$ fold change +1.31 and +1.63), is structurally similar to the copper-binding protein CopM (PDB 5ffc; $E = 4 \times 10^{-10}$, 73% query coverage, 43% identity). Over the same days the CopR iModulon increased in the mixed co-culture (raw $p = 0.09$ and $0.06$; **Table S4**), and its highest-weighted gene is a second DUF305 protein encoded next to *copR* on plasmid pANL. SYNPCC7942_RS05895 is not a member of the CopR iModulon, and both observations are weak; together they point to a possible copper-related response that remains to be tested.

#### Supplementary figures

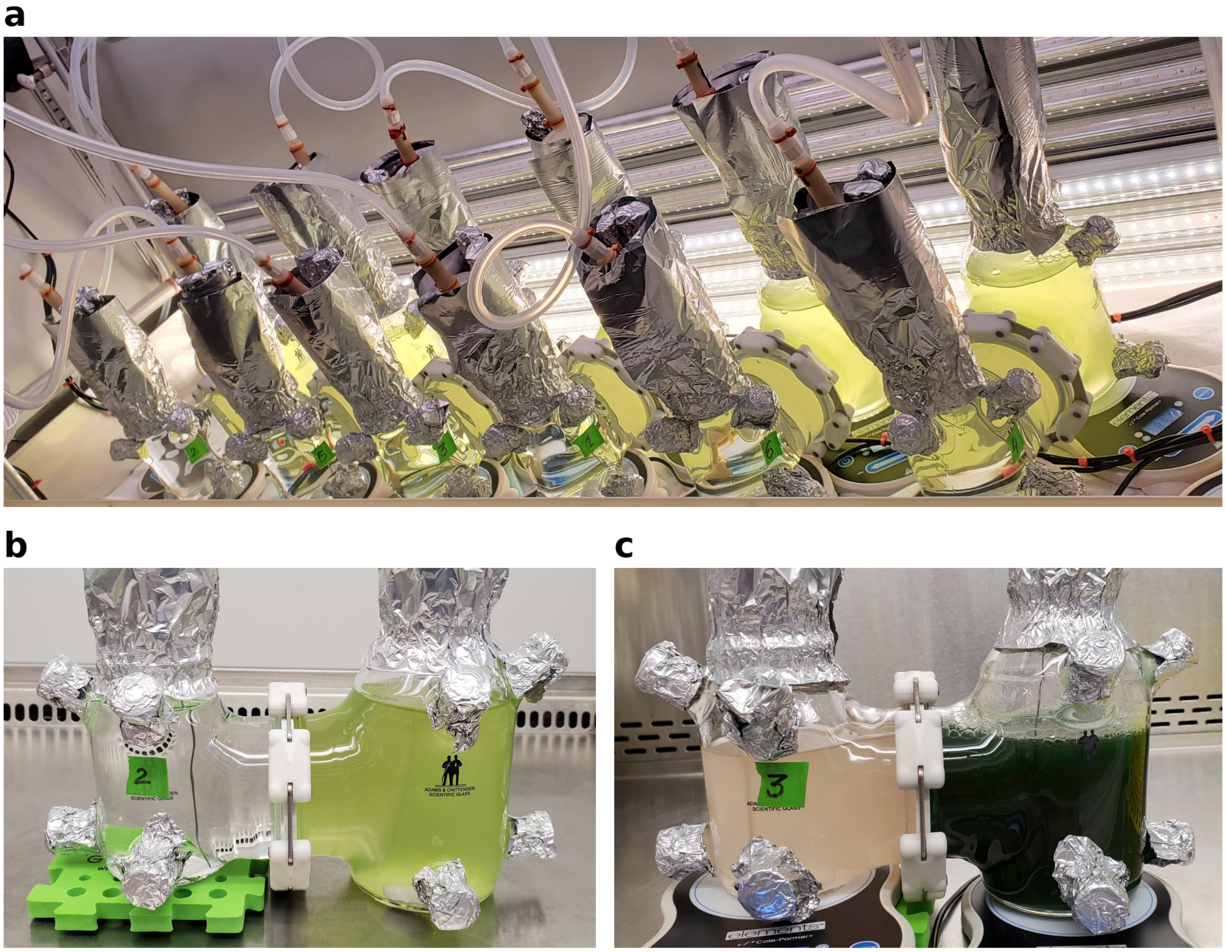

**Figure S1. Two-chamber membrane bioreactors (mBR) at the start of the experiment and early in co-culture. (a)** The six mBRs at time 0, on magnetic stir plates under LED panels, before *R. toruloides* was added; reactors 1 to 3 are co-cultures and reactors 4 to 6 are axenic controls. **(b)** Reactor 2 at time 0: the chamber that later received *R. toruloides* holds cell-free medium, and the *S. elongatus* chamber is dilute. **(c)** Reactor 3 on day 6: *R. toruloides* is growing but still dilute, and the *S. elongatus* chamber is dense. The two chambers of each mBR are joined at an NW63 flange that holds a 0.22 µm PVDF membrane.

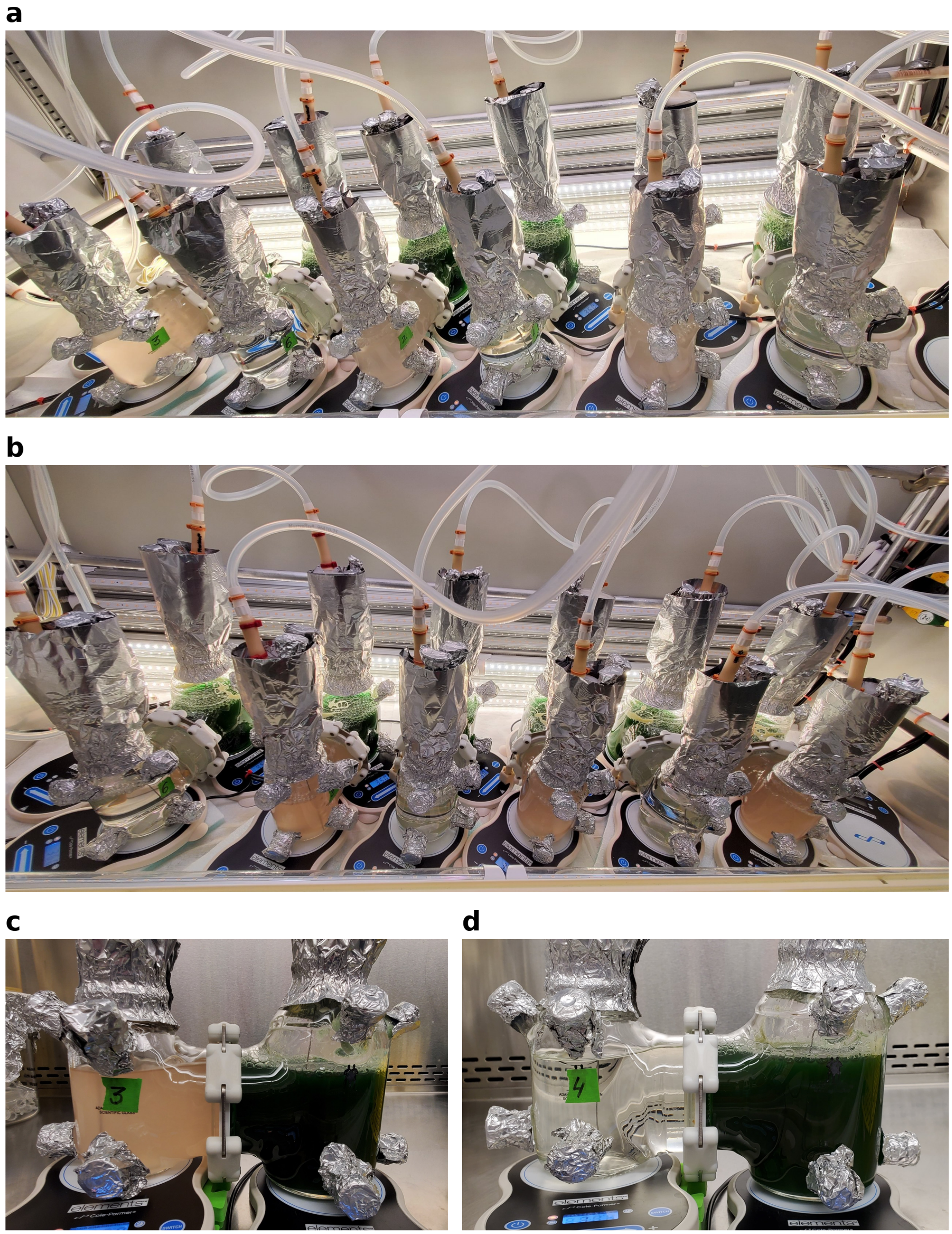

**Figure S2. Membrane bioreactors later in co-culture. (a)** Days 5 to 6: *R. toruloides* is growing but still dilute in the co-culture reactors, while the cell-free chambers of the axenic controls stay clear. **(b)** Days 8 to 9: the *R. toruloides* chambers are denser, and the cell-free chambers stay clear. **(c)** Reactor 3 (co-culture) on days 8 to 9: dense *R. toruloides* (tan) beside dense *S. elongatus* (green). **(d)** Reactor 4 (axenic control) on days 8 to 9: cell-free medium (clear) beside very dense *S. elongatus*.

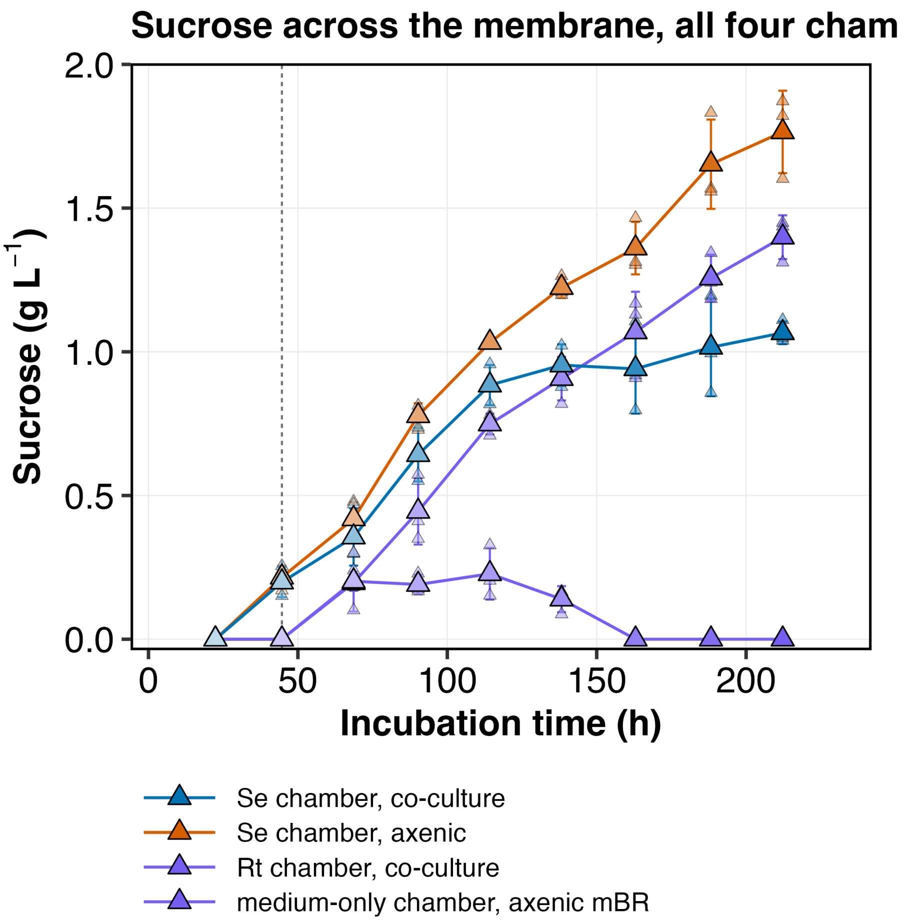

**Figure S3. Sucrose in the four chambers of the two membrane bioreactors.** Se chamber and Rt chamber of the co-culture mBR, and Se chamber and blank (medium only) chamber of the axenic mBR. The blank chamber shows sucrose crossing the membrane with no consumer, reaching 1.40 g $L^{-1}$ by day 9, while the *R. toruloides* chamber never exceeds 0.23 g $L^{-1}$ and is below the assay limit from day 7. Points are replicates (pale fill) and group means (full fill), n = 3; error bars are one standard deviation. Every series also appears in **Figure 1b to d**. Values: **Table S1**.

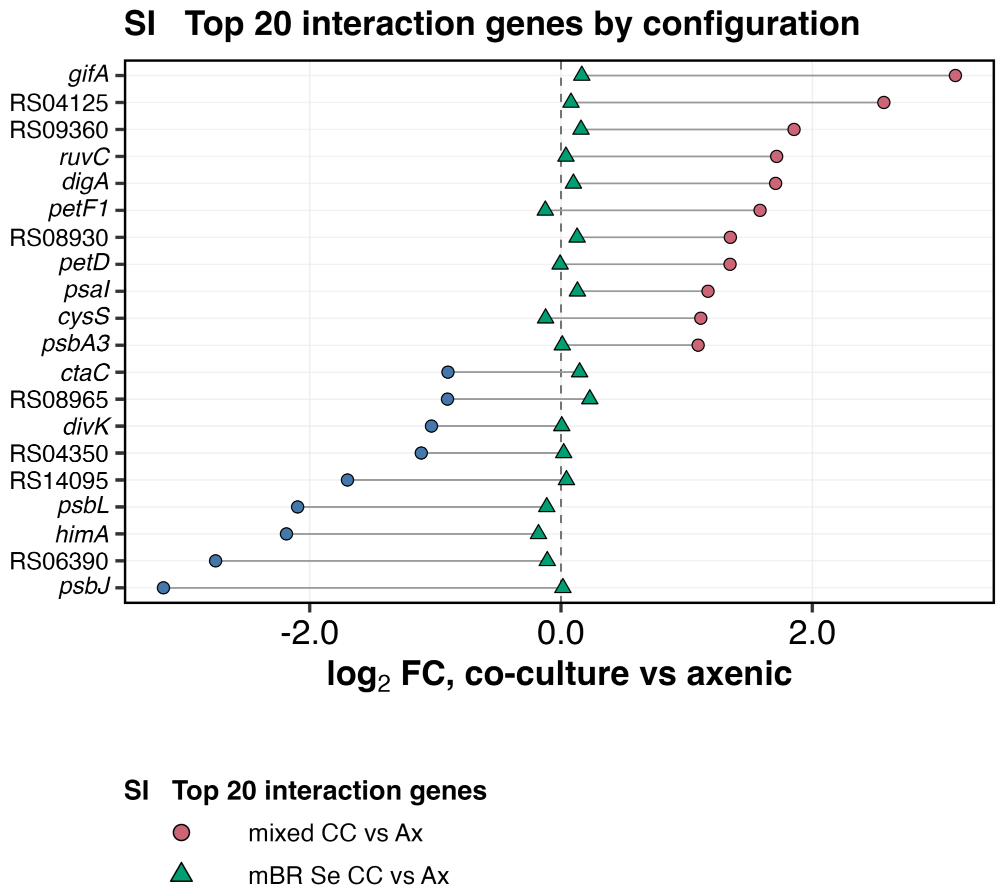

**Figure S4. The 20 interaction genes with the smallest p-adjusted.** Each row is one gene from the configuration by culture interaction (**Figure 4b**): $\log_2$ fold change, co-culture minus axenic, in the mixed reactor (circles; red, higher; blue, lower) and in the mBR Se chamber (teal triangles), joined by a line. The mBR Se chamber values sit near zero. Genes are labelled by gene name where one exists, otherwise by RefSeq locus tag. Values: **Table S3**, sheet Top20_interaction.

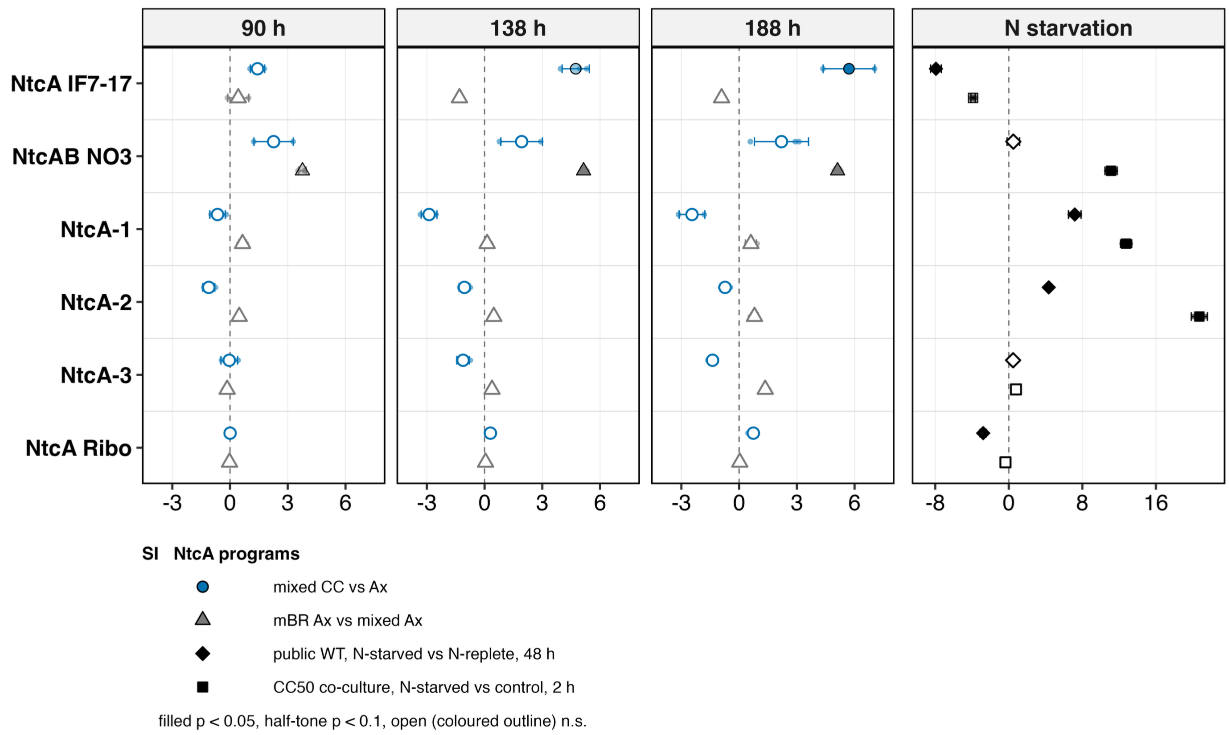

**Figure S5. NtcA programs against two nitrogen-starvation references.** The six NtcA iModulons as activity differences from their own controls by sampling day (90, 138 and 188 h): co-culture (mixed CC vs Ax) and reactor configuration (mBR Ax vs mixed Ax). The right column, on its own scale, shows two nitrogen-starvation references computed with the same null: wild-type *S. elongatus* 48 h without nitrogen minus nitrogen-replete (public data), and the CC50 co-culture 2 h without nitrogen minus with nitrogen (this laboratory). Mean and one standard deviation; marks give the DiMA test (* $p < 0.05$, ** $p < 0.01$, *** $p < 0.001$, † $p < 0.1$). Values: **Table S4**, sheets DiMA and N_starvation_references.

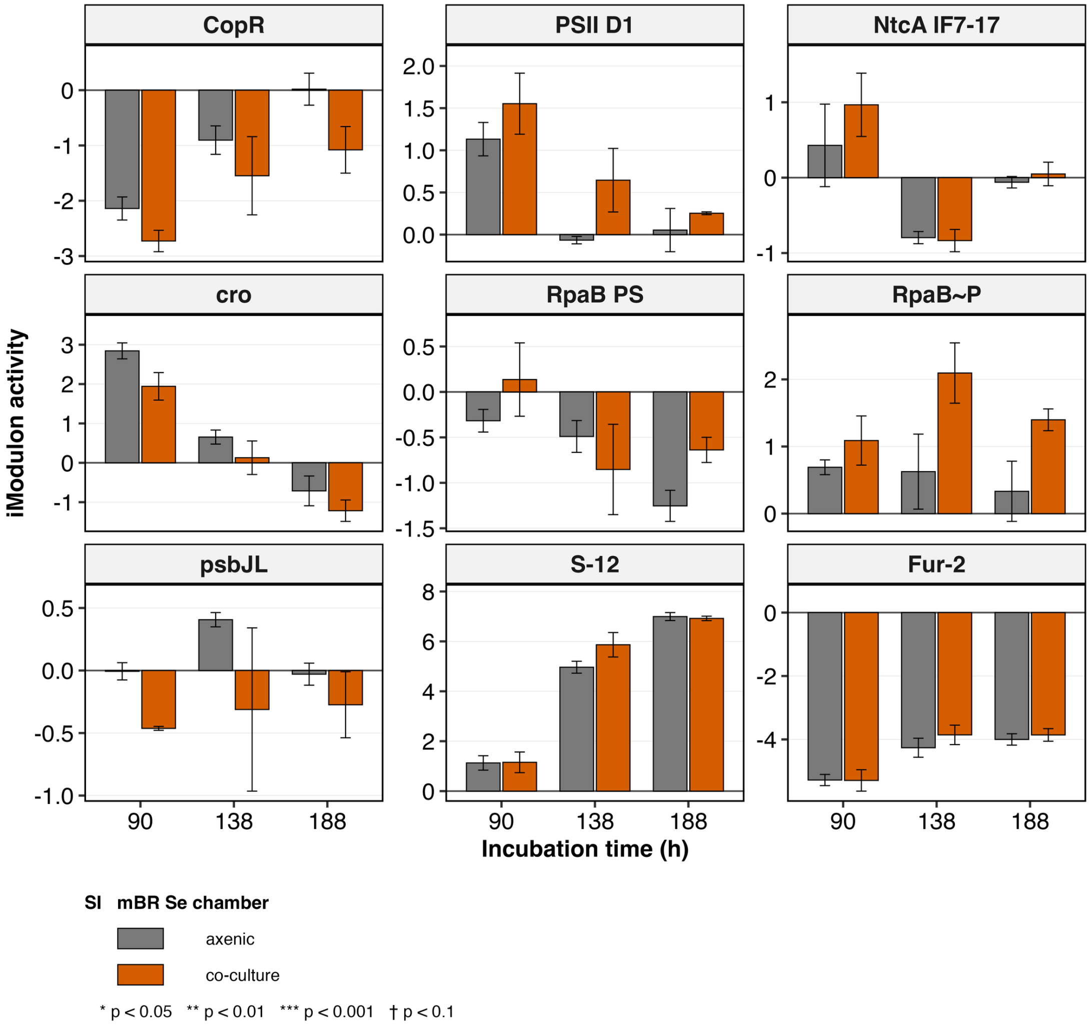

**Figure S6. iModulon activity in the mBR Se chamber.** Activity of the nine iModulons of **Figure 5** in the mBR Se chamber, axenic and co-culture, by sampling day, on the same anchor as **Figure 5b** (mean of the matched day-4 mixed axenic samples, Se_ax_90_1 to 3, subtracted per iModulon). Bars are group means, error bars one standard deviation; marks as in **Figure 5b**. No iModulon reaches $p < 0.05$ on any day. Values: **Table S4**, sheets DiMA and Activities.

#### Supplementary tables

Each **Table Sx** is an Excel workbook with the complete data. The compact summary sheets are reproduced here for reading; the workbook remains the authoritative source.

**Table S1. Growth of *S. elongatus* and *R. toruloides* in the mixed reactor and the membrane bioreactor.** The workbook Table_S1_growth.xlsx holds the time base; specific growth rates by two methods; *S. elongatus* co-culture over axenic density by day in each configuration and pooled over days 3 to 8 at reactor level; mBR axenic over mixed axenic density; the decomposition of **Fig. 1e**; *R. toruloides* mBR over mixed density; and curve fits. Two summary sheets are reproduced here. (a) Pooled co-culture effect: geometric mean ratio of co-culture to axenic density with the Welch 95% confidence interval (CI) and p value, computed at reactor level on ln density (each reactor averaged over days 3 to 8; n = 3 reactors per group). (b) Ratio of mBR co-culture to mixed axenic density on each sampling day, separated into a reactor-type part (mBR axenic over mixed axenic) and a partner part (mBR co-culture over mBR axenic); the total is the product of the two parts. CC, co-culture; Ax, axenic control; mBR, membrane bioreactor; Se, *S. elongatus*.

*(a) Pooled co-culture effect on* S. elongatus *density, days 3 to 8 (sheet Pooled_days3to8)*

| Comparison | Window | Ratio | 95% CI | p |
|---|---|---|---|---|
| Mixed reactor, CC / Ax | Days 3 to 8 | 1.092 | 1.051 to 1.135 | 0.003 |
| mBR Se chamber, CC / Ax | Days 3 to 8 | 0.986 | 0.923 to 1.052 | 0.500 |

*(b) Decomposition of* ***Fig. 1e*** *(sheet Decomposition)*

| Day | Total: mBR CC / mixed Ax | Reactor-type part: mBR Ax / mixed Ax | Partner part: mBR CC / mBR Ax |
|---|---|---|---|
| Day 4 | 1.061 | 1.083 | 0.980 |
| Day 6 | 1.103 | 1.102 | 1.001 |
| Day 8 | 1.089 | 1.105 | 0.985 |
| Day 10 | 1.050 | 1.075 | 0.977 |

**Table S2. Exometabolome of the mixed reactor and the membrane bioreactor.** The workbook Table_S2_exometabolome.xlsx holds the compound annotation and identification markers; $\log_{10}$ fold changes and q values for all 98 compounds in the four comparisons; the direction summary; Se-chamber increases that are also higher in the Rt chamber than in cell-free medium; normalization sensitivity; divergence by day; total ion signal; non-detects; and citrate by day. The direction summary is reproduced here. Fold changes are the first group minus the second on $\log_{10}(x + 1)$ intensity, from a linear model with group and day as terms across days 1 to 9; q is the Benjamini-Hochberg adjusted p across the 98 compounds. Wilcoxon p is a signed-rank test of the 98 fold changes against zero. CC, co-culture; Ax, axenic control; blank, cell-free chamber of the axenic mBR; FC, fold change; mBR, membrane bioreactor; Se, *S. elongatus*; Rt, *R. toruloides*.

*(a) Direction of change by comparison (sheet Direction_summary)*

| Comparison | Median $\log_{10}$ FC | Compounds below zero | Wilcoxon p | Lower / higher at $q < 0.05$ |
|---|---|---|---|---|
| Mixed reactor, CC vs Ax | -0.131 | 76% | $4.10 \times 10^{-7}$ | 46 / 9 |
| mBR Se chamber, CC vs Ax | 0.129 | 22% | $2.90 \times 10^{-8}$ | 2 / 46 |
| mBR Rt chamber, CC vs blank | 0.111 | 32% | 0.006 | 13 / 41 |
| mBR Ax vs mixed Ax | -0.048 | 56% | 0.580 | 36 / 32 |

**Table S3. Gene-level transcriptome of *S. elongatus*.** The workbook Table_S3_transcriptome.xlsx holds differentially expressed gene counts by comparison and day; all differentially expressed and interaction genes with fold changes and adjusted p; sensitivity to repeated sampling and normalization; limma against DESeq2; regulon, gene set, KEGG and COG enrichment; sigma factors; the top 20 interaction genes; quality-control evidence for the excluded sample; and the re-annotation of genes without an assigned function. Two summary sheets are reproduced here. (a) Gene counts from limma-voom with reactor identity as a random effect, for 35 samples from days 4, 6 and 8: genes at adjusted $p < 0.05$, and differentially expressed genes (DEGs; adjusted $p < 0.05$ and an absolute $\log_2$ fold change above 1) that are higher or lower in the first-named group. The interaction is the configuration by culture term, the mixed co-culture effect minus the mBR Se chamber co-culture effect. Days averaged uses all three sampling days. (b) The 20 interaction genes with the smallest adjusted p, ranked; fold changes are co-culture minus axenic. Locus tags and old locus tags are shown without their prefixes. Labels are gene names where one exists, otherwise the locus tag without its prefix. CC, co-culture; Ax, axenic control; mBR, membrane bioreactor; Se, *S. elongatus*.

*(a) Differentially expressed gene counts (sheet DEG_counts)*

| Comparison and sampling day | Genes at $q < 0.05$ | DEGs ($q < 0.05$, $\lvert\log_2FC\rvert > 1$) | DEGs higher | DEGs lower |
|---|---|---|---|---|
| *Mixed CC vs Ax* | | | | |
| Days averaged | 1468 | 51 | 35 | 16 |
| Day 4 | 331 | 20 | 10 | 10 |
| Day 6 | 1230 | 91 | 56 | 35 |
| Day 8 | 1269 | 127 | 90 | 37 |
| *mBR Se CC vs Ax* | | | | |
| Days averaged | 71 | 1 | 1 | 0 |
| Day 4 | 0 | 0 | 0 | 0 |
| Day 6 | 246 | 1 | 1 | 0 |
| Day 8 | 0 | 0 | 0 | 0 |
| *Interaction (mixed minus mBR Se)* | | | | |
| Days averaged | 1087 | 55 | 30 | 25 |
| Day 4 | 120 | 18 | 8 | 10 |
| Day 6 | 1047 | 138 | 80 | 58 |
| Day 8 | 812 | 112 | 73 | 39 |
| *mBR Ax vs mixed Ax* | | | | |
| Days averaged | 1468 | 40 | 14 | 26 |
| Day 4 | 1175 | 49 | 17 | 32 |
| Day 6 | 1232 | 55 | 16 | 39 |
| Day 8 | 1021 | 61 | 33 | 28 |
| *Mixed CC vs mBR Se CC* | | | | |
| Days averaged | 1747 | 65 | 39 | 26 |

*(b) The 20 interaction genes with the smallest adjusted p (sheet Top20_interaction)*

| Rank | Locus tag (SYNPCC7942_) | Old locus tag (Synpcc7942_) | Label | $log_2FC$, mixed CC vs Ax | $log_2FC$, mBR Se CC vs Ax | $p_{adj}$, interaction |
|---|---|---|---|---|---|---|
| 1 | RS13745 | 1174 | *psbJ* | -3.164 | 0.015 | $2.46 \times 10^{-20}$ |
| 2 | RS04125 | 0797 | RS04125 | 2.569 | 0.079 | $5.77 \times 10^{-15}$ |
| 3 | RS04350 | 0841 | RS04350 | -1.112 | 0.022 | $4.53 \times 10^{-14}$ |
| 4 | RS14095 | – | RS14095 | -1.699 | 0.043 | $6.68 \times 10^{-14}$ |
| 5 | RS11420 | 2248 | *himA* | -2.185 | -0.179 | $7.77 \times 10^{-14}$ |
| 6 | RS08475 | 1662 | *cysS* | 1.113 | -0.122 | $1.03 \times 10^{-12}$ |
| 7 | RS04130 | 0798 | *ruvC* | 1.716 | 0.039 | $1.45 \times 10^{-11}$ |
| 8 | RS04640 | 0900 | *gifA* | 3.139 | 0.166 | $1.96 \times 10^{-11}$ |
| 9 | RS06020 | 1175 | *psbL* | -2.096 | -0.113 | $5.76 \times 10^{-11}$ |
| 10 | RS08620 | 1693 | *divK* | -1.031 | 0.006 | $9.31 \times 10^{-11}$ |
| 11 | RS08930 | 1757 | RS08930 | 1.348 | 0.128 | $1.23 \times 10^{-10}$ |
| 12 | RS13255 | 2602 | *ctaC* | -0.900 | 0.148 | $1.96 \times 10^{-10}$ |
| 13 | RS11905 | 2343 | *psaI* | 1.170 | 0.131 | $2.27 \times 10^{-10}$ |
| 14 | RS04600 | 0893 | *psbA3* | 1.092 | 0.010 | $2.85 \times 10^{-10}$ |
| 15 | RS09360 | 1845 | RS09360 | 1.855 | 0.159 | $2.85 \times 10^{-10}$ |
| 16 | RS08470 | 1661 | *digA* | 1.708 | 0.099 | $5.15 \times 10^{-10}$ |
| 17 | RS07660 | 1499 | *petF1* | 1.584 | -0.125 | $5.21 \times 10^{-10}$ |
| 18 | RS06390 | 1247 | RS06390 | -2.749 | -0.109 | $6.44 \times 10^{-10}$ |
| 19 | RS08965 | 1764 | RS08965 | -0.903 | 0.229 | $8.18 \times 10^{-10}$ |
| 20 | RS11855 | 2332 | *petD* | 1.345 | -0.007 | $8.18 \times 10^{-10}$ |

**Table S4. iModulon activity of *S. elongatus*.** The workbook Table_S4_iModulons.xlsx holds differential iModulon activity (DiMA) for all 78 iModulons in three comparisons on three sampling days; activities of every sample, raw and on the matched day-4 anchor; two nitrogen-starvation references; and the 24 CC50 perturbations against their common control. All sheets are large matrices and are provided in the workbook only.
